# Outcome contingency selectively affects the neural coding of outcomes but not of tasks

**DOI:** 10.1101/375642

**Authors:** David Wisniewski, Birte Forstmann, Marcel Brass

## Abstract

Value-based decision-making is ubiquitous in every-day life, and critically depends on the contingency between choices and their outcomes. Only if outcomes are contingent on our choices can we make meaningful value-based decisions. Here, we investigate the effect of outcome contingency on the neural coding of rewards and tasks. Participants performed a reversal-learning task in which reward outcomes were contingent on trial-by-trial choices, and performed a ‘free choice’ task in which rewards were random and not contingent on choices. We hypothesized that contingent outcomes enhance the neural coding of rewards and tasks, which was tested using multivariate pattern analysis of fMRI data. Reward outcomes were encoded in a large network including the striatum, dmPFC and parietal cortex, and these representations were indeed amplified for contingent rewards. Tasks were encoded in the dmPFC at the time of decision-making, and in parietal cortex in a subsequence maintenance phase. We found no evidence for contingency-dependent modulations of task signals, demonstrating highly similar coding across contingency conditions. Our findings suggest selective effects of contingency on reward coding only, and further highlight the role of dmPFC and parietal cortex in value-based decision-making, as these were the only regions strongly involved in both reward and task coding.

## Introduction

Making decisions is an integral part of our life. Most of these choices are value-based, i.e. they are made with expected outcomes in mind. Value-based choices are made in separate stages: we first evaluate all options, and then select the option with the highest subjective value ^1^. After implementing the chosen behavior ^2^, predicted and experienced outcomes are compared, and prediction errors are computed ^3–5^. This dopamine-mediated learning signal ^6^ indicates the need to update our internal models of action-outcome contingencies, which then leads to an adaption of future behavior.

This process is modulated by various properties of choice outcomes, e.g. their magnitude ^7^. However, one crucial aspect has received little attention in the past: to which degree our choices directly control possible outcomes. Clearly, whether or not we believe our choices to directly *cause* their outcomes affects decision-making considerably. If we know that a specific behavior predictably leads to a desired outcome, we will choose it more often ^8^. If we know that our behavior and desired outcomes are only weakly correlated, or not correlated at all, we might not prioritize any specific behavior. Here, we focus on investigating the effects of high vs low control of the outcomes of one’s own choices.

In principle, varying degrees of control of choice outcomes can affect two key processes: outcome valuation and the implementation of chosen behavior. Some previous research in non-human primates ^9^, and humans ^10, 11^ demonstrated that choice-contingent outcomes are processed differently than non-contingent outcomes. Importantly, one might expect similar effects on neural representations of the chosen behavior that is operational for receiving the reward as well. One might expect, for example, chosen behaviors to be shielded more strongly against interference if outcomes are contingent on them ^12^, as not performing the behavior as intended is potentially costly. For non-contingent outcomes the need for shielding is lower, as e.g. executing the wrong behavior has no effect on received outcomes (see ^13^ for a related argument, but ^14^). Previous work demonstrated that implementation of chosen actions, is supported by a brain network including the frontopolar ^15^, lateral prefrontal and parietal cortex ^16–18^. Some initial evidence suggests that rewarding correct performance indeed enhances neural task representations ^19^, but this work did not address the issue of varying degrees of control over choice outcomes.

Here, we report an experiment investigating the effects of control over choice outcomes on value-based decision making. We used a value-based decision task and multivariate pattern analysis methods (MVPA ^20^) to assess the effects of reward contingency (choice-contingent vs. non-contingent rewards) on valuation and, more importantly, on choice implementation. We first hypothesized that reward contingency affects the neural coding of outcomes ^9, 11^. We further assessed whether implementation of chosen behavior (i.e. coding of chosen tasks) is similarly affected by contingency. We hypothesized that the lateral prefrontal cortex, and especially the parietal cortex to play a key role in the implementation of chosen behavior. The parietal cortex represents chosen tasks and actions ^1, 17^, subjective stimulus and action values ^21, 22^, as well as associations between choice options and their outcomes ^23^. Here, we tested whether task representations in these brain regions were enhanced when rewards were choice-contingent vs when they were not.

## Results

Participants in this experiment implemented two different sets of stimulus-response (SR) mappings (labelled task X and task Y, Figure 1). In half of the trials, they performed a probabilistic reward reversal-learning task (contingent reward (CR), see ^24^), choosing between performing two different tasks that led to either high or low reward outcomes in each trial. Crucially, these reward outcomes were contingent on the specific choices made in each trial, and contingencies changed across the course of the experiment. In the other half of the trials, participants performed a ‘free choice’ task (non-contingent reward, NCR). After each trial, they again received high or low rewards, but the outcomes were uncorrelated with the choices and participants had no means of controlling the reward outcomes in these trials.

**Figure 1.**
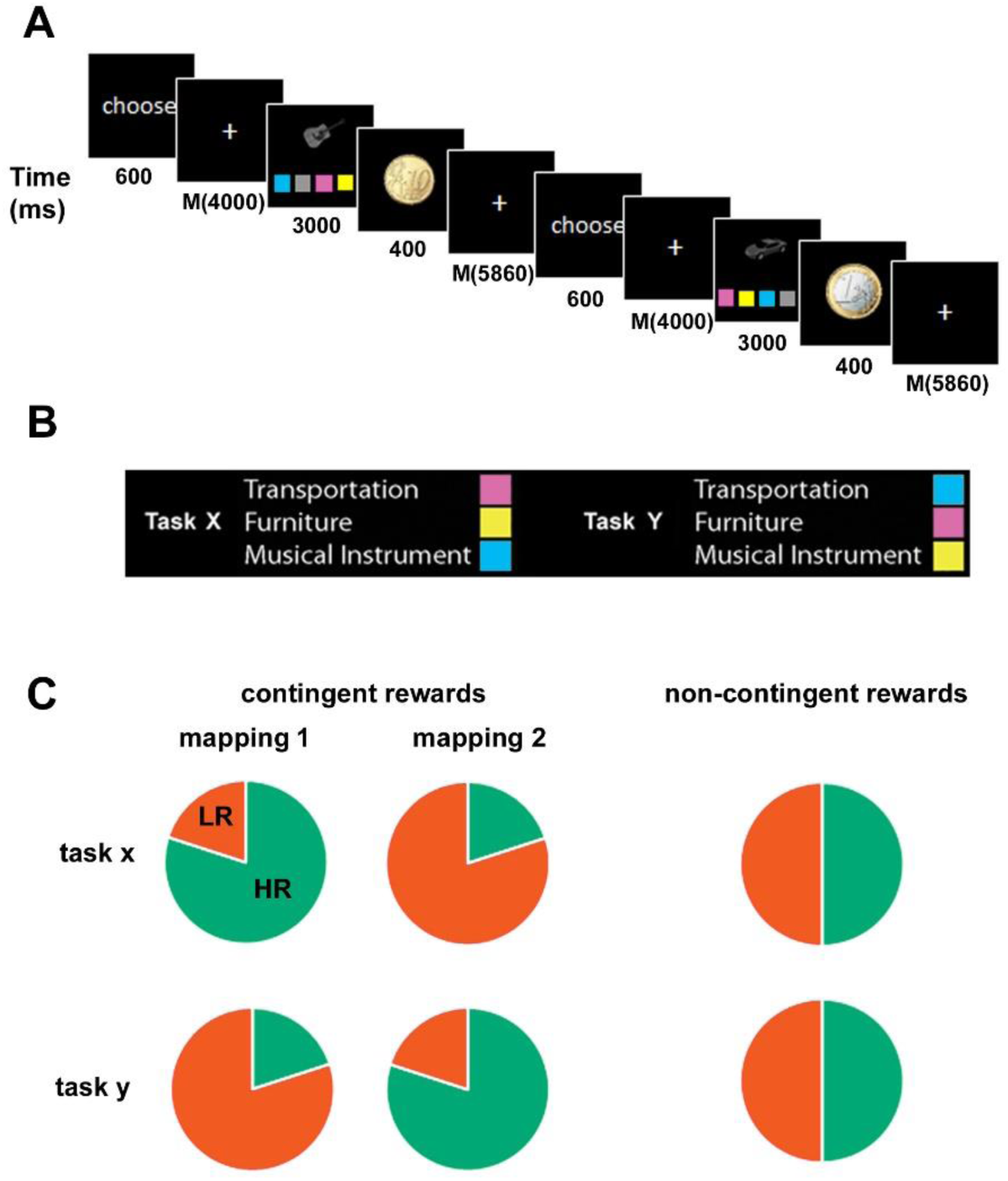
Experimental paradigm. **A.** Trial structure. Each trial started with the cue ‘choose’ presented on screen, indicating that subjects should decide which task to perform on that trial. After a variable delay, the task screen was presented for a fixed duration. Reward feedback was presented subsequently after each trial (high reward = 1€, low reward = 10€cents, no reward). All trials were separated by variable inter trial intervals. **B.** Tasks. Subjects were instructed to identify the visual object presented on screen randomly drawn out of 9 objects, belonging to 3 categories (means of transportation, furniture, musical instruments). Each category was associated with a colored button, and subjects were instructed to press the corresponding button. Two different sets of stimulus-response mappings were learned, and labelled task x and task y. On each trial, subjects had the free choice which of the two tasks to implement. **C.** Reward conditions. In contingent trials, subjects performed a probabilistic reversal-learning task. In each trial one of the two tasks yielded a high reward with a high probability (80%), and a low reward with a low probability (20%). The other task had the opposite reward contingencies. Which task yielded higher rewards depended on the current task-reward-mapping, which changed across the experiment. In non-contingent trials, subjects also received high and low reward outcomes, which were assigned randomly (50%/50%) and were not contingent on specific task choices.

### Behavioral results

We first assessed whether error rates or reaction times (RT) differed between the two sets of SR mappings (labelled task X, Y), or the reward contingency condition (CR, NCR). The average error rate across all subjects was 5.89% (SEM = 0.74%). Thus, subjects were able to perform the task accurately. There was no evidence for an effect of reward condition on error rates (Bayes Factor (BF10) = 0.88). Error trials were removed from all further analyses. A repeated-measures ANOVA on RTs including the factors SR mapping and contingency revealed evidence for the absence of any RT differences between the two contingency conditions (BF10 = 0.01, Figure 2A). We further found evidence for the absence of RT differences between the specific SR-mappings implemented in each trial (Figure 1, BF10 = 0.01), showing that both SR-mappings were equally difficult to perform. There was moderate evidence for the absence of an interaction between task and reward contingency (BF10 = 0.23).

**Figure 2.**
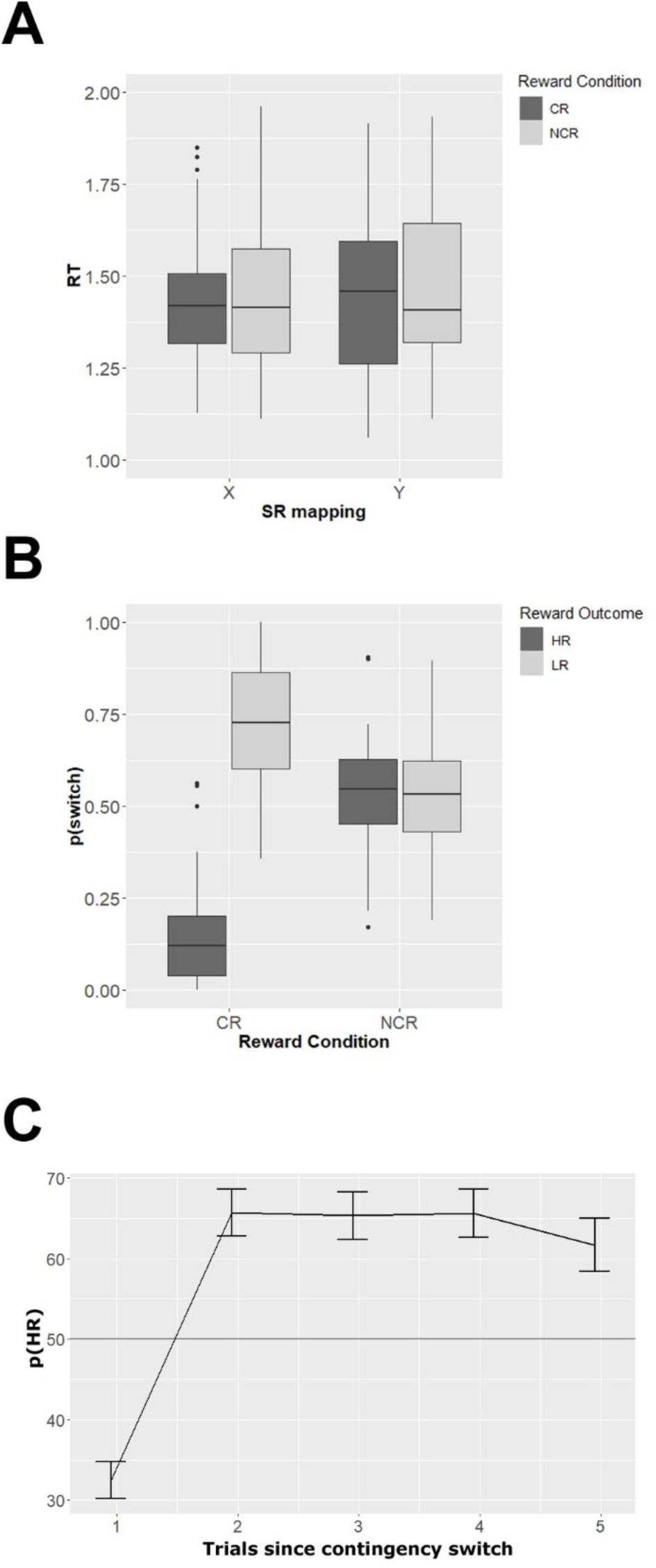
Behavioral Results. **A.** Reaction Times (RT). The box plots depict reaction times for each combination of stimulus-response mapping and reward condition. Contingent (CR) trials are shown in dark grey, non-contingent (NCR) trials are shown in light grey. **B.** Switch probabilities. Probability to switch away from the current task as a function of previous reward (high = HR dark grey, low = LR light grey), separately for contingent (CR) and non-contingent (NCR) trials. **C.** Probability to choose the high reward task in CR trials (pHR), as a function of how many trials passed since the last reward contingency switch. Participants chose below chance (50%) on trials immediately following a contingency switch (‘perseveration’), and then quickly switched to choosing the HR task on subsequent trials. All error bars depict the SEM.

We then assessed whether subjects showed choice biases towards one of the two SR-mappings, which might indicate stable preferences and in turn affect fMRI analyses (see below). We found no choice bias in CR trials, 50.35% (SEM = 1.49%), which did not differ from 50%, BF10 = 0.18, BF10_r=1.41_ = 0.09). The same was true for NCR trials, 51.40% (SEM = 1.75%), BF10 = 0.24, BF10_r=1.41_ = 0.12, indicating that subjects did not exhibit strong preferences for specific SR mappings. Next, we assessed performance in the reversal learning (CR) task. For this purpose, we computed how often subjects chose the highly reward (HR) task, and this value should be above 50% if they succeeded in learning the changing reward contingencies. Subjects chose the HR task in 61.10% (SEM = 1.74%) of the CR trials, which was above 50% (BF10 >150). As a control analysis, we computed the number of HR outcomes in NCR trials, which should be 50% if choices were uncorrelated with outcomes. This was indeed the case, HR outcomes = 49.47% (SEM = 0.84%), t-test vs 50% (BF10 = 0.21). Importantly, we found CR trials to lead to HR outcomes more often than NCR trials (56.4%, SEM = 1.15%, BF10 > 150), demonstrating that subjects succeeded in choosing tasks strategically in CR trials to maximize their reward outcomes. In order to assess how they maximized their outcomes, we computed the probability to switch to a different task just after receiving a high or a low reward (p_switchHR_, p_switchLR_). As expected, we found subjects to stay in the task that led to a high reward on the previous trial, p_switchHR_ = 15.71% (SEM = 2.60%), and switch away from the task that led to a low reward on the previous trial, p_switchLR_ = 71.08% (SEM = 3.20%, Figure 2B). We found no such difference in NCR trials, p_switchHR_ = 53.12% (SEM = 2.83%), p_switchLR_ = 52.85% (SEM = 2.46%). An ANOVA with the factors reward contingency (CR, NCR) and reward outcome (HR, LR) identified a main effect of reward outcome on p_switch_ (BF10 > 150), no main effect of contingency (BF10 = 1.85), and a strong interaction (BF10 > 150). This demonstrates that subjects employed a win-stay loose-switch strategy and had clear reward-expectations selectively in CR trials, and that reward outcomes in NCR trials had no immediate effect on task choices. Next, we assessed learning of changing reward contingencies in CR trials, by computing the probability to choose the highly rewarded task (p_HR_), as a function of trials passed since the last contingency change. We expected subjects to systematically choose the LR task immediately following such a change (‘perseveration’), but then quickly learn about the new contingency mapping. Our results confirmed these expectations (Figure 2C). While subjects rarely picked the HR task immediately following a contingency switch (p_HR_ = 32.48%, SEM = 2.28%), this value increased dramatically already on the subsequent trial (p_HR_ = 65.65%, SEM = 2.88%, paired t-test, BF10 > 150). There was no evidence for changes in p_HR_ on the following four trials, all BF_10_s < 0.36, demonstrating that learning occurred mostly within the first trial following a contingency switch. Lastly, we hypothesized that subjects stayed in the same task longer in CR trials, as compared to NCR trials, and found this to be the case (Supplementary Analysis 1).

### Multivariate decoding of reward outcome values

#### Baseline analysis

We first set out to determine whether outcome contingency affects the neural coding of outcome magnitude. For this purpose, we identified brain regions encoding outcome values (high vs low) at the time of feedback presentation (*baseline decoding*, collapsing across CR and NCR trials). We found an extensive network to encode outcome values including striatal subcortical brain regions, as well as large parts of the prefrontal and parietal cortex (Figure 3A, mean accuracy = 60.52%, SEM = 1.11%). This contrast does not *only* capture specific reward value signals, it might also reflect effects caused by differences in reward outcomes, like attention or motor preparation. In order to address at least some of these confounding factors, we regressed out RTs out of the data before performing MVPA ^25^, but found no effect strong effects on our results (Supplementary Figure 1).

**Figure 3:**
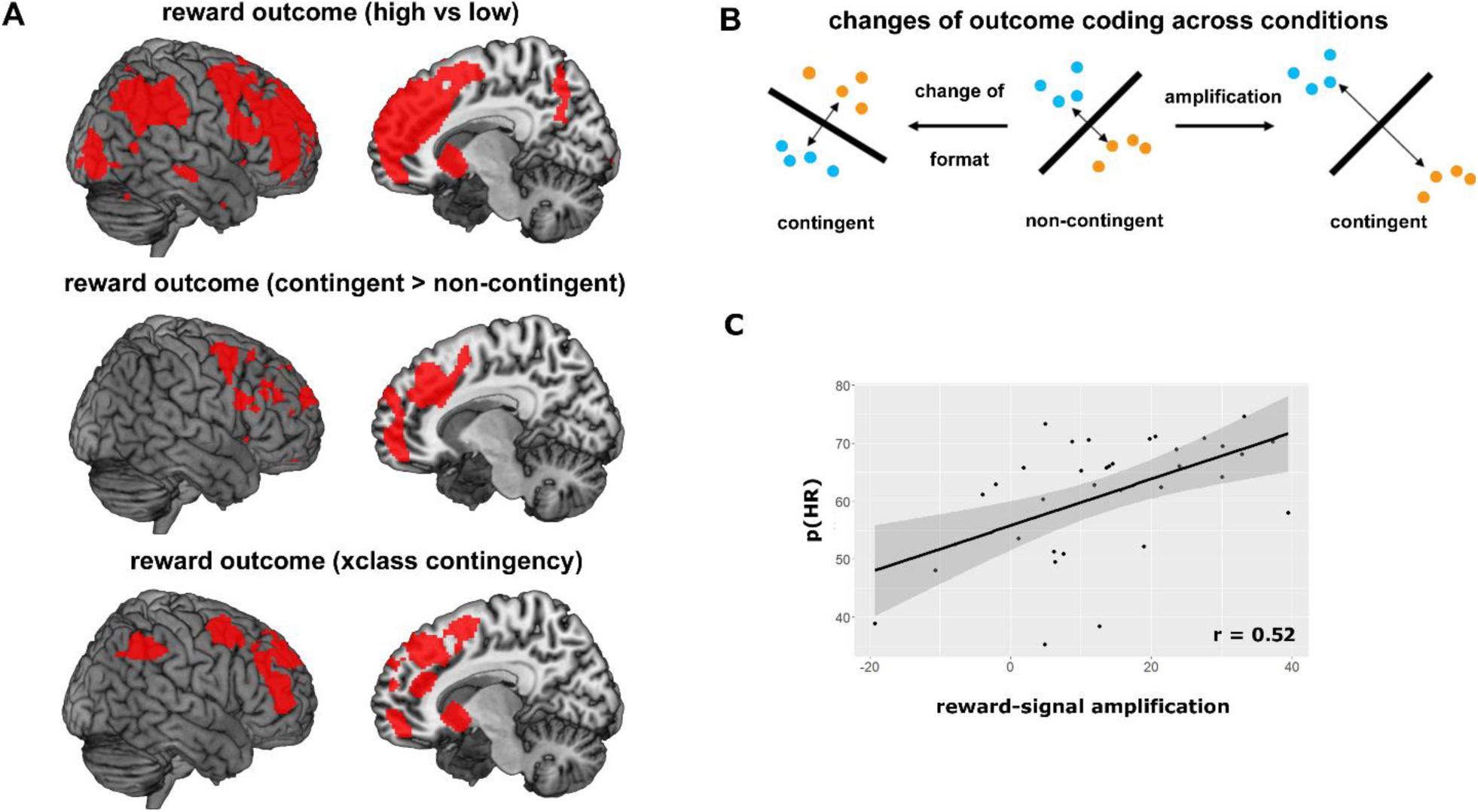
Reward-related brain activity. **A.** Multivariate decoding of reward outcome value. Above: baseline decoding. Depicted are regions that encoded the value of reward outcomes (high vs. low, combined across CR and NCR conditions). The regions identified were used as masks for the following analyses. Results are displayed at p < 0.05 (FWE corrected). Middle: regions showing stronger outcome coding in contingent (CR) than in non-contingent (NCR) trials. Below: regions encoding reward values in similar formats in both contingency conditions, as tested using a cross-classification (xclass) analysis. **B.** Amplification vs change of format of neural coding. Most regions identified in A showed both stronger decoding in CR trials, and similar formats across both contingency conditions. This is compatible with an amplification or gain increase of neural codes. In the middle, a hypothetical example of a pattern decoding is depicted. High reward trials are depicted as blue, low reward trials as orange dots. The classifier fits a decision boundary to separate the two distributions. If this code changes between the two contingency conditions (left), decoding is still possible at similar accuracy levels as before, but a classifier trained on NCR trials will be unsuccessful in classifying CR trials. If this code is amplified in the CR condition however (right), the same patterns become more easily separable. The same classifier will be will be successful in both conditions and accuracies will increase. See ^27^ for more information. **C.** Correlation of reward signal amplification and successful performance. This plot shows the correlation between the degree of reward signal amplification (accuracy in CR trials – accuracy in NCR trials), and successful performance in CR trials (probability to choose the high reward task, p(HR)). Each dot is one subject, and the line depicts a fitted linear function with 95% confidence intervals (gray area).

#### Differences in outcome coding

Subsequently, we assessed whether these outcome signals were modulated by reward contingency, hypothesizing that contingent rewards showed stronger decoding results than non-contingent rewards. We repeated the same decoding analysis separately for CR and NCR trials and assessed whether any region from the baseline analysis showed stronger outcome coding in CR (mean accuracy = 64.17%, SEM = 2.01%) than in NCR trials (mean accuracy = 54.32%, SEM = 1.24%, using a within-subjects ANOVA and small-volume correction, p < 0.001 uncorrected, p < 0.05 FWE corrected). We found the striatum, bilateral lateral PFC, dACC, anterior medial PFC, and IPS to show stronger reward outcome coding for contingent rewards. The opposite contrast (NCR > CR) yielded no significant results (p < 0.001 uncorrected, p < 0.05 FWE corrected). This effect cannot be explained be differences in outcome value per se, as the reward magnitude did not differ across the conditions, only contingency did.

#### Similarities in outcome coding

In a last step, we assessed whether any of the brain regions identified in the baseline analysis showed similar coding across contingency conditions, using a cross-classification approach (see Materials and Methods for more details, ^26^). We trained a classifier on CR trials, and tested it on NCR trials, and vice versa, and found the striatum, lateral and medial PFC, dACC, and IPS to encode rewards similarly across conditions (mean accuracy = 55.52%, SEM = 1.10%). This pattern of results suggests that the neural code for different reward outcomes did not change across contingency conditions, yet outcome signals were still stronger in CR than in NCR trials. This points to an amplification or gain-increase of reward-related signals through contingency ^27^. Interestingly, post-hoc analyses revealed that the difference in decoding accuracies between CR and NCR trials (i.e. the strength of the signal-amplification), correlated positively with successful performance in CR trials (Figure 3C), r = 0.52, 95%CI = [0.26, 0.76] (for more details see Supplementary Analysis 2). The more subjects amplified their contingent reward-representations, the more they succeeded in performing the reversal-learning task, which links outcome coding to behavior.

### Multivariate decoding of tasks

#### Maintenance period

##### Baseline analysis

For the task coding, we expected to see similar effects as for the reward outcome signals. In a first analysis, we focused on the task maintenance period (from ‘choose’ cue onset to task screen onset, see Materials and Methods for more details, and ^28^ for a similar approach). During this time, subjects maintain the abstract SR mapping they want to implement, without yet being able to implement specific motor actions. As in the outcome decoding analysis, we first combine CR and NCR trials to identify all regions encoding tasks (*baseline* decoding, see also ^16^). We found two brain regions to contain task information, the left posterior parietal cortex (mean accuracy = 54.61%, SEM = 0.65%), spanning over the midline into the right parietal cortex, and the right anterior middle frontal gyrus (aMFG, mean accuracy = 54.66%, SEM = 0.89%, see Figure 4A, Table 1). The parietal cluster identified here partly overlapped with the parietal cluster identified in the outcome decoding analysis, suggesting parietal involvement in both reward and task processing.

**Figure 4:**
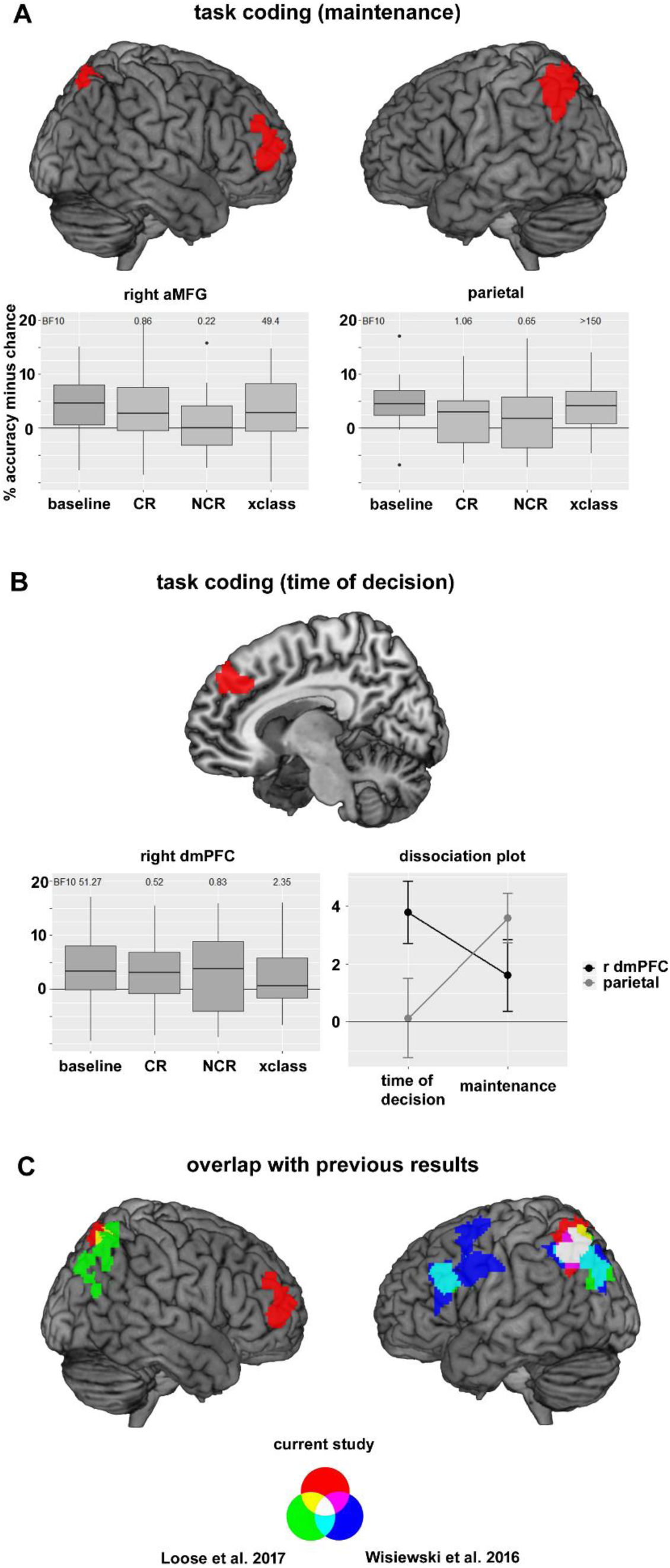
Task coding. **A.** Task coding during maintenance. Results from the baseline decoding analysis are depicted above. Two clusters passed the significance threshold (p < 0.001 uncorrected at the voxel level, p < 0.05 FWE corrected at the cluster level), one in the parietal cortex, and one in the right anterior MFG. Accuracies were then extracted for the contingent (CR), non-contingent (NCR), and contingency cross-classification (xclass) task decoding analyses. Results can be seen in the boxplots. Above the plots, Bayes factors (BF10) of a t-test vs. chance level are shown. BF10 for the baseline analysis is not reported, as this analysis was used to define the ROIs, and running additional statistical tests on this data would constitute double dipping. **B.** Task coding at the time of decision-making. Above, the dmPFC ROI used in this analysis (from ^29^) is depicted. The box plot depicts results from our data in this ROI, for all four analyses performed (baseline, CR, NCR, xclass). The dissociation plot depicts a double dissociation between two ROIs (right dmPFC, as defined using data from ^29^, and the left parietal cortex, as defined using data from ^17^), and two time points in the trial (time of decision-making, maintenance). All error bars represent SEM. **C.** Overlap with previous results. Results from the current study (red) are overlain on previous findings from ^17^ (blue), and ^16^ (green). All results are based on task decoding analyses (searchlight decoding, radius = 3 voxels, C = 1, chance level = 50%), albeit with different specific tasks being contrasted in each study. Despite this fact, all three studies find task information around the intraparietal sulcus. Findings in the PFC are less consistent.

**Table 1:** Baseline task decoding

|  |  |  |  | MNI coordinates (peak) |  |  |
| --- | --- | --- | --- | --- | --- | --- |
| Brain region | Side | Cluster size | Mean accuracy (SEM) | X | Y | Z |
| parietal cortex | Bilateral | 2427 | 54.61% (0.65%) | -10 | -60 | 60 |
| anterior MFG | Right | 955 | 54.66% (0.89%) | 32 | 58 | 18 |

Results are shown for a statistical threshold of $p < 0.001$ (uncorrected) at the voxel level and $p < 0.05$ (FWE corrected) at the cluster level. Chance level is 50%.

##### Differences in task coding

If contingent outcomes indeed enhance task coding, we should see higher accuracies in CR, than in NCR trials. To test this, we repeated the task decoding analysis separately for CR and NCR trials, and extracted accuracy values in the two ROIs found in the baseline task decoding analysis. We found no task information in the parietal cortex in these two analyses (CR: 51.29%, SEM = 0.91%, BF10 = 1.06; NCR: 51.73%, SEM = 1.44%, BF10 = 0.64), found no evidence for stronger task coding in CR than in NCR trials (BF10 = 0.16), and no evidence for stronger task coding in NCR than in CR trials (BF10 = 0.19). A similar pattern of results was found in the right aMFG (CR: 51.79%, SEM = 1.37%, BF10 = 0.85; NCR: 50.48%, SEM = 1.35%, BF10 = 0.22; CR > NCR: BF10 = 0.40; NCR > CR: BF10 = 0.10). Thus, we find no evidence for an effect of reward contingency on task representations, despite the fact that behavior clearly differed between the two reward conditions, and that contingency has been found to modulate the coding of reward outcomes. As these two analyses are based on only half as many trials as the baseline analysis, a reduction in statistical power could explain the absence of any differences. In order to test this, we performed an additional control analysis, in which we showed that task coding is modulated by outcome magnitude (but not by outcome contingency), demonstrating that modulatory effects can be found in principle in this data-set with the same statistical power (Supplementary Analysis 3).

##### Similarities in task coding

As in the outcome coding analysis, we also tested whether task representations were similar across the two contingency conditions, using a cross-classification approach. We found both the parietal cortex (54.03%, SEM = 0.76%, BF10 > 150), as well as the right aMFG (53.71%, SEM = 1.16%, BF10 = 49.39) to show similar task coding across contingency conditions. We also tested whether results from this cross-classification differed from the baseline accuracies, finding moderate evidence for an absence of any differences (parietal cortex BF10 = 0.23, aMFG BF10 = 0.25). These results show that the parietal cortex and aMFG encode tasks using a general format that is similar across reward-contingency conditions.

#### Time of decision-making

Our experimental design allows us to independently assess task-related signals both at the time of decision-making and the subsequent maintenance period as these were separated by a jittered delay period (see Materials and Methods for more details). We thus repeated the same task decoding analyses on the time of decision-making. For this purpose, we used the right dmPFC cluster found previously ^29^ as a ROI, which was found to encode task information at the time of decision-making. Despite the differences in experimental design, we found task information in the baseline analysis (53.76%, SEM = 1.07%, BF10 = 51.27, Figure 4B) here as well. Accuracies were not above chance in CR trials (51.95%, SEM = 1.83%, BF10 = 0.52) and NCR trials (52.45%, SEM = 1.71%, BF10 = 0.83), and did not differ from each other (BF10 = 0.15). We found anecdotal evidence for successful task cross-classification in this region (52.03%, SEM = 0.98%, BF10 = 2.35), although the baseline and xclass analyses still did not differ (BF10 = 1.64). Interestingly, the dmPFC cluster partly overlapped with results from the reward outcome decoding. Additionally, we found a double dissociation in task coding between the right dmPFC and left parietal cortex (Figure 4B, both ROIs defined a priori from previous experiments ^17, 29^), with the former only encoding tasks at the time of decision-making, and the latter only encoding tasks during intention maintenance. An ANOVA using the factors ‘time in trial’ (time of decision vs intention maintenance) and ‘ROI’ (right dmPFC vs left parietal cortex) revealed moderate evidence for a time × ROI interaction (BF10 = 5.39). Furthermore, the right dmPFC only encoded tasks at the time of decision (BF10 = 51.27), but not during intention maintenance (BF10 = 0.68). Conversely, the left parietal cortex only encoded tasks during intention maintenance (BF10 > 150), but not at the time of decision (BF10 = 0.19). This suggests a temporal order of task processing in the brain, with task information first being encoded in dmPFC, and then in parietal cortex (but see ^30^). Additionally, a post-hoc analysis revealed that decoding accuracies in the dmPFC at the time of decision-making correlated positively with trait impulsivity (Supplementary Analysis 2).

#### Control analyses

In order to test the generalizability of our results, we repeated the same analyses on several a-priori ROIs, taken from two previous experiments ^16, 17^, which tested for effects of cognitive control, and free choice on task coding (during a maintenance period), respectively. Overall, we were able to replicate the above effects in these a-priori defined ROIs, although findings were much more consistent in parietal than in prefrontal brain regions (Figure 4C, Supplementary Analysis 4). In a similar vein, we also assessed task information in the multiple demand network ^31, 32^, and were able to replicate our findings in the parietal cortex (see Supplementary Analysis 5 for more information).

It has been shown previously that RTs might affect task decoding results, leading to false-positive findings ^25^. Although others failed to replicate this effect ^33^, and we found no RT effects on the group level, we decided to conservatively control for RT effects on the single-subject level nonetheless,. After regressing RT-related effects out of the data for each subject, we still found the parietal cortex to encode tasks (54.61%, SEM = 0.65%, BF10 > 150), and also found the task coding in the cross-classification analysis (54.03%, SEM = 0.76%, BF10 > 150). The same was true for the aMFG (54.66%, SEM = 0.89%, BF10 > 150; and 53.71%, SEM = 1.16%, BF10 = 23.38 respectively). Results in the baseline and xclass analysis were equal in both regions, BFs10 <= 0.30. These results thus mirror the main analysis above, showing that RT-related variance cannot explain task decoding results in our experiment.

In order to validate the decoding procedure, we also extracted task decoding accuracies from a region not involved in performing this task, the primary auditory cortex. As expected, we found accuracies not to differ from chance level in this region (49.64%, SEM = 0.93%, BF10 = 0.13), showing that the task decoding analysis was not biased towards positive accuracy values and specific to the regions reported.

Lastly, we performed two control analyses to directly assess possible biases in our decoding procedure (Supplementary Analysis 6), and effects of error rates and/or choice biases on our task decoding results (Supplementary Analysis 7). None of the factors was found to affect our results.

## Discussion

Here, we investigated whether controlling reward outcomes modulates the neural coding of either outcomes or tasks. Subjects performed a probabilistic reward reversal learning task, in which outcomes were contingent on specific choices. They also performed a free choice task with non-contingent reward outcomes, in which outcomes were not under their direct control. Although we found reward contingency to modulate outcome valuation, contrary to our expectations we found no effects on choice implementation. Furthermore, we found two main brain regions to be crucial for encoding tasks and reward outcomes: the right dmPFC and the left parietal cortex (around the IPS). The dmPFC was found to encode chosen tasks at the time of decision-making, and simultaneously encoded reward outcome values, emphasizing its role in both value-related with intentional control processes. While the parietal cortex encoded reward outcomes at the time of decision-making, it encoded chosen tasks during a subsequent maintenance phase. We found a double dissociation between both regions, with the dmPFC encoding tasks only at the time of decision-making, and the parietal cortex only during intention maintenance.

Much previous research on the effects of reward motivation on cognition investigated the effects of reward prospect ^34^. These findings demonstrated that positive reinforcement improves cognition, as compared to no reinforcement at all. However, an equally important and often overlooked property of reinforcement is the degree of control we have in reaching it. Sometimes, an action will cause outcomes in a fairly clear way (e.g. hitting a light switch), other times, that link will be less close (e.g. refreshing your Facebook timeline). Previous work has shown that the strength of such action-outcome contingencies modulates the neural processing of reward outcomes ^11^, and here we directly demonstrate how neural coding changes and how this change affects behavior. Outcomes (and correlated processes) are encoded more strongly if they are contingent on choices, as compared to when they are not. Their representational format further does not change strongly across contingency conditions. This is compatible with an amplification of outcome representations through contingency (Figure 3B), where representations do not change but become more separable in neural state space (see ^13^ for a similar argument). This is in line with predictions from gain-theories of motivation, which suggest that subcortical dopaminergic neurons can modulate their gain ^6^, making them more or less sensitive to changes in rewards (see also ^35^). Here, we demonstrate such gain increases in subcortical dopaminergic regions and beyond. This effect is unlikely to reflect simple motor processes, as regressing RTs out of the data did not alter reward decoding results. It might be related to reward processing, but given the current data we cannot fully exclude that other related (e.g. attentional) processes contribute as well. Crucially, the strength of this gain increase was correlated with successful performance, the more subjects increased neural gain, the more successful they were in performing the reversal-learning task. Thus, our data demonstrates how reward-related signals changed, and that they are behaviorally relevant for successful performance.

Importantly, in order for this value signal to lead to actual rewards, chosen behavior has to be implemented as intended first ^36^. One might thus expect contingency to lead to stronger task shielding and coding ^12^, as the costs of confusing the two highly similar tasks are potentially high. However, we found no evidence for such effects. On the contrary, we found evidence for a similar coding of tasks across both contingency conditions. This finding informs current debates on the nature of task coding in the brain ^27^. On the one hand, some have argued for flexible task coding especially in the fronto-parietal cortex ^32, 37^, often based on the multiple-demand network theory ^31^. This account predicts that task coding should be stronger when task demands are high ^32^, or when correct performance is rewarded ^19^. Despite our efforts to replicate these findings in our data-set, we found no evidence for an influence of reward contingency on task coding. This was despite the fact that behavior differed between these conditions and that value-related signals were affected by reward contingency. One might argue that our analysis had insufficient statistical power to detect true effects, but a control analysis revealed that we can in principle detect task coding differences in this data-set, making this explanation unlikely.

On the other hand, others have argued that the same task representations could be used in multiple different situations (i.e. ‘multiplexing’ of task information), and that this allows us to flexibly react to novel and changing demands ^14^. Multiplexing predicts that task information should be invariant across different contexts, which has been shown previously ^16–18^. Here, we extend these findings, by showing that tasks are encoded in a format that is similar across contingency conditions in both frontal and parietal brain regions, strengthening the idea of multiplexing of task information in the brain. One possible alternative explanation for this finding might be that subjects were highly trained in performing the two tasks, and were at their performance ceiling. This might make a modulation of task coding too small to detect. Although we cannot fully exclude this interpretation, we want to point out that contingency did have robust effects on behavior. Also, most related previous experiments trained their subjects, those that found modulatory effects ^19, 32^ and those that did not ^17^. We thus believe this alternative explanation to be unlikely. Overall, our task decoding results are in line with the idea of multiplexing of task information in the brain. Future research will have to test more systematically which environmental conditions lead to multiplexing of task information in the brain, and which do not.

One key region identified here was the dmPFC. It is supports effort-based foraging choices ^29^, and here we show its involvement in a reward reversal learning task. The dmPFC is important for cognitive control, supporting rule and action selection ^38^, and processing uncertainty ^39^. It has further been associated with valuation processes, anticipating positive outcomes ^40^, and encoding reward prediction errors ^41^. In this experiment, we demonstrated that the dmPFC is specifically involved in encoding tasks only at the time at which a choice is made, other regions later maintain that choice outcome until it can be executed. We also demonstrated the dmPFC to encode outcome values at the same time. Please note that we do not claim this value signal to only represent the magnitude of reward outcomes, it might also represent related processes (e.g. attention). Nevertheless, the cause of this effect are different outcome values, and this highlights the importance of dmPFC in linking valuation to strategic decision-making, suggesting how it might support goal-directed behavior ^42^.

The second key region identified in this experiment was the left parietal cortex (IPS). The parietal cortex is a key region for cognitive control ^43^, and is part of the multiple demand network ^31^, a set of brain regions characterized by their high flexibility to adapt to changing demands. Previous work on non-human primates demonstrated that the prefrontal cortex flexibly switches between representing different control-related information within single trials ^44^. Our results show that the parietal cortex in humans exhibits similar flexibility ^23^, it encodes both control-related and value-related variables. This provides further evidence for its flexibility, and it will be interesting to assess how the parietal cortex links value-related and control-related variables in future experiments. Given its involvement in foraging behavior ^22^, the previous choice and outcome history might affects current choice representations in this brain region. Future studies optimized to investigate this question will help shedding more light on this issue.

In sum, we assessed whether controlling outcomes affects outcome and task processing in the brain. By comparing choices that are informed by expected outcomes as well as choices that are not, we linked largely parallel research on ‘free choice’ and value-based decision-making. While we found strong effects on outcome processing, we found no such effects on choice implementation. Our results further highlight the importance of both the dmPFC and parietal cortex in bridging valuation and executive processes in the brain.

## Materials and Methods

### Participants

A total of 42 subjects participated in this experiment (20 males, 21 females, 1 other). The average age was 22.6 years (min = 18, max = 33 years), 41 subjects were right-handed, one was left-handed. All subjects had normal or corrected-to-normal vision and volunteered to participate. Subjects gave written informed consent and received between 45€ and 55€ for their participation. The experiment was approved by the local ethics committee. Seven subjects showed excessive head movement in the MR scanner (>4mm) and were excluded.

### Experimental Design

#### Trial structure

The experiment was programmed using PsychoPy (version 1.85.2, psychopy.org, RRID:SCR_006571 ^45^). Each trial started with the presentation of a fixation cross centrally on-screen (300ms, Figure 1A). This was followed by the presentation of a choice cue (‘CHOOSE’, 600ms), which instructed subjects to freely choose one of the two tasks to perform in this trial. After a variable delay period (2000-6000ms, mean duration = 4000ms), the task screen was presented for a total of 3000ms, irrespective or RT. Similar to ^29^, the task screen consisted of a visual object presented centrally on screen (Figure 1B). This object was picked pseudo-randomly out of a pool of 9 different objects in 3 categories: musical instruments, furniture, means of transportation. Below, 4 colored squares were presented (magenta, yellow, cyan, gray), with the square positions being mapped onto 4 buttons, operated using the left and right index and middle fingers. Subjects were given the option to choose which of two sets of stimulus-response (SR) mappings to apply to the presented object (e.g. labelled task ‘X’ = means of transportation → magenta, furniture → yellow, and musical instruments → cyan. task ‘Y’ = means of transportation → cyan, furniture → magenta, and musical instruments → yellow, the grey button was never task-relevant and included merely to balance left and right button presses). Thus, one button was correct for each task in each trial, and subjects were instructed to react as quickly and accurately as possible. Choices were inferred from the pressed buttons. We use the term ‘task’ here to describe a specific link between stimuli and responses, and we do not claim that different cognitive processes are required to perform each task. Importantly, the position of the colored buttons on screen was pseudo-randomized in each trial, preventing preparation of specific motor responses before the onset of the task screen. Furthermore, which set of S-R-mappings was called task ‘X’ and task ‘Y’ was counter-balanced across subjects. After the task screen offset, a reward feedback was presented (image of 1€ coin (high reward, HR), 10€cent coin (low reward, LR), or red circle (no reward), 400ms). After a variable inter-trial-interval (4000-14000ms, geometrically distributed, mean duration = 5860ms), the next trial began.

#### Reward conditions

Subjects were rewarded for correct performance on every trial. There were a total of two different reward conditions: contingent rewards (CR) and non-contingent rewards (NCR). In the NCR condition, the chosen reward in each trial was determined randomly. Irrespective of the chosen task, subjects had a 50% chance of receiving a high and a 50% chance of receiving a low reward (Figure 1C). Subjects were instructed to choose tasks randomly in this condition, by imagining flipping a coin in their head on each trial ^18^. In the CR condition, subjects performed a probabilistic reward reversal-learning task, similar to ^24^. In each trial, one task led to a high reward with an 80% and a low reward with a 20% probability (HR task). These probabilities were reversed for the other task (LR task). Subjects were merely instructed that reward contingencies were stable across a few trials, but could change over time, and needed to infer the current contingency from the trial-by-trial reward feedback. After choosing the HR task for 3 consecutive trials, reward contingencies reversed without warning with a chance of 50% in each subsequent trial (similar to ^24^). At the end of the experiment, 15 trials were chosen randomly (both from CR and NCR trials), and whichever reward was earned in these trials was paid out as a bonus payment to the subjects. This ensured that subjects were motivated in each trial and condition.

This reward manipulation was designed to vary the degree of control over choice outcomes. Choices in CR trials directly affected reward outcomes, which made expected outcomes a highly relevant task feature to take into account during decision-making. Choices in NCR trials were unrelated to reward outcomes, and expected outcomes were irrelevant for decision-making. We ensured that the reward condition was uncorrelated to all other design variables (target stimulus, delay duration, button mapping, ITI duration), and previous trial conditions were not predictive of current trial conditions. This ensured that all trials were IID, and estimated neural signals were not confounded.

#### Procedure and Design

Subjects first performed a training session which lasted about 1h10min 1-5 days before the MR session. They learned to perform the task and completed 3 runs of the full experiment. This minimized learning effects during the MR session, which can be detrimental to MVPA. In the MR session, subjects performed 5 identical runs of this experiment, with 60 trials each. Each run contained 2 blocks with CR and 2 blocks with NCR trials. The length of each block was between 10 and 14 trials, and all trials were all separated by a long and variable ITI. CR and NCR blocks alternated and block order was counterbalanced across runs for each subject. Each block started with either ‘Contingent block now starting’ or ‘Non-contingent block now starting’ presented on screen (5000ms). This mixed blocked and event-related design minimized cross-talk and interference between the reward conditions. Reward contingencies did not carry over from a one to the next CR block, in order to make each block independent from previous performance. Each run also contained 20% (n=12) catch trials. In these trials, subjects were externally cued which task to perform, and the delay between cue and task execution was only 1000ms. Catch trials were included to prevent subjects from choosing all tasks in a block at its beginning. For instance, in an NCR block, subjects could theoretically decide upon a whole sequence of tasks at the beginning of that block (e.g. X,X,X,Y,X,Y,Y,X,…), and then only implement that fixed sequence in each trial. In order to encourage subjects to make a conscious choice in each individual trial, catch trials were included to frequently disrupt any planned sequence of task choices, making such a strategy less feasible. To maximize the salience of catch trials, correct performance always led to a high reward. Catch trials were excluded from all analyses.

#### Additional measures

After completing the MR session, subjects filled in multiple questionnaires. They answered custom questions (e.g., How believable were the instructions? Was one task more difficult than the other?) and the following questionnaires: behavioral inhibition / activation scale (BISBAS ^46^), need for cognition (NFC ^47^), sensitivity to reward / punishment (SPSRQS ^48^), and impulsivity (BIS11 ^49^). We also acquired pupil dilation data while subjects performed the experiment in the MR scanner. Pupil dilation data is not the focus of the current paper, and is not reported.

### Image acquisition

fMRI data was collected using a 3T Magnetom Trio MRI scanner system (Siemens Medical Systems, Erlangen, Germany), with a standard thirty-two-channel radio-frequency head coil. A 3D high-resolution anatomical image of the whole brain was acquired for co-registration and normalization of the functional images, using a T1-weighted MPRAGE sequence (TR = 2250 ms, TE = 4.18 ms, TI = 900 ms, acquisition matrix = 256 × 256, FOV = 256 mm, flip angle = 9°, voxel size = 1 × 1 × 1 mm). Furthermore, a field map was acquired for each participant, in order to correct for magnetic field inhomogeneities (TR = 400 ms, TE_1_ = 5.19 ms, TE_2_ = 7.65 ms, image matrix = 64 × 64, FOV = 192 mm, flip angle = 60°, slice thickness = 3 mm, voxel size = 3 × 3 × 3 mm, distance factor = 20%, 33 slices). Whole brain functional images were collected using a T2*-weighted EPI sequence (TR = 2000 ms, TE = 30 ms, image matrix = 64 × 64, FOV = 192 mm, flip angle = 78°, slice thickness = 3 mm, voxel size = 3 × 3 × 3 mm, distance factor = 20%, 33 slices). Slices were orientated along the AC-PC line for each subject.

### Statistical Analysis

#### Data Analysis: Behavior

All behavioral analyses were performed in R (RStudio version 1.1.383, RRID:SCR_000432, www.rstudio.com). We first characterized subjects’ performance by computing error rates and reaction times (RT). We tested for potential effects of reward condition on error rates using a Bayesian two-sided paired t-tests (using *ttestBF* from the BayesFactor package in R). Error trials (trials with wrong button presses, or with RTs <300ms) were then removed from the data analysis. In order to identify potential effects of the different SR mappings and reward contingency condition on RTs, we performed a Bayesian repeated measures ANOVA (using *anovaBF* from the BayesFactor package in R). This ANOVA included the factors SR mapping and reward contingency, and outputs Bayes Factors (BF) for all main effects and interaction terms. We did not expect to find RT differences between SR mappings, but did expect RTs to be lower in the CR condition, as compared to the NCR condition. All Bayesian tests were performed using the default prior (Cauchy prior, r=.707). We performed additional robustness checks using different priors (r=1, r=1.41), which did not change our results in most cases. When priors did affect the interpretation of results, these results are reported (e.g. BF10_r=1.41_), otherwise we only report results using the default prior.

The Bayesian hypothesis testing employed here allows quantifying the evidence in favor of the alternative hypothesis (BF10) *and* the null hypothesis, allowing us to conclude whether we find evidence for or against a hypothesized effect, or whether the current evidence remains inconclusive ^50^. We considered BFs between 1 and 0.3 as anecdotal evidence, BFs between 0.3 and 0.1 as moderate evidence, and BFs smaller than 0.1 as strong evidence against a hypothesis. BFs between 1 and 3 were considered as anecdotal evidence, BFs between 3 and 10 as moderate evidence, and BFs larger than 10 as strong evidence for a hypothesis.

Given that subjects were free to choose between the two tasks, some subjects might have shown biases to choosing one of the two SR mappings more often (although that would not have led to a higher overall reward, if anything biases should lower overall rewards). In order to quantify biases, we computed the proportion of trials in which subjects chose each SR mapping, separately for the CR and NCR conditions, and tested whether this value differed from 50% using a two-sided Bayesian t-test. The output BF was interpreted in the same way as in the previous analysis.

Choices in CR trials were assessed by quantifying how well subjects performed the probabilistic reversal learning task. If subjects were reliably able to determine which of the two tasks was currently the HR task, they should have chosen that task more often than expected by chance (50%). Thus the proportion of HR task choices (p_HR_) in CR trials is our main measure of how successful subjects were in performing the task. This measure was compared to chance level using a one-sided Bayesian t-test. We also computed this measure as a function of trials that passed since the last contingency switch. We expected subjects to systematically choose the LR task immediately following such a change (‘perseveration’), but then quickly learn about the new contingency mapping. We tested whether p_HR_ changed across trials by using Bayesian paired t-tests. Furthermore, we assessed the earned reward outcomes, expecting p_HR_ to be higher in CR, than in NCR trials (where it should be 50%). This was tested using a paired one-sided Bayesian t-test.

In order to describe the strategies employed to maximize reward outcomes, we computed the probability to switch to a different task immediately following a high or a low reward (ps_witchHR_, p_switchLR_). We expected subjects to follow a “win stay loose switch” strategy (WSLS), staying in a highly rewarded task and switching away from a lowly rewarded task. To test this, we performed a two-factorial Bayesian ANOVA, including the factors reward outcome (HR, LR) and reward contingency (CR, NCR). We expected to see a main effect of outcome on p_switch_, and an interaction, as WSLS should be specifically applied to CR trials only.

#### Data Analysis: fMRI

fMRI data analysis was performed using Matlab (version R2014b 8.4.0, RRID:SCR_001622, The MathWorks) and SPM12 (RRID:SCR_007037, www.fil.ion.ucl.ac.uk/spm/software/spm12/). Raw data was imported according to BIDS standards (RRID:SCR_016124, http://bids.neuroimaging.io/), and then was unwarped, realigned and slice time corrected.

##### Multivariate decoding of reward outcomes

In a first step, we assessed whether we can replicate previous findings demonstrating contingency effects on reward processing ^10, 11^. For this purpose, we estimated a GLM ^51^ for each subject. For each of the 5 runs we added regressors for each combination of reward value (HR vs LR) and contingency (CR vs NCR). All regressors were locked to the reward feedback onset, the duration was set to 0. Regressors were convolved with a canonical haemodynamic response function (as implemented in SPM12). Estimated movement parameters were added as regressors of non-interest to this and all other GLMs reported here.

###### Baseline decoding

In a next step, we performed a decoding analysis on the parameter estimates of the GLM. A linear support-vector classifier (SVC ^52, 53^), as implemented in *The Decoding Toolbox* ^54^, was used using a fixed regularization parameter (C = 1). We performed searchlight decoding ^20, 28^, which looks for information in local spatial patterns in the brain and makes no a prior assumptions about informative brain regions. Searchlight radius was set to 3 voxels, and we employed run-wise cross-validation. We contrasted HR trials (from both CR and NCR trials) with LR trials (again from CR and NCR trials). The resulting accuracy maps were normalized to a standard space (Montreal Neurological Institute template as implemented in SPM12), and smoothed (Gaussian kernel, FWHM = 6mm) in order to account for potential differences in information localization across subjects. Group analyses were performed on the accuracy maps using voxel-by-voxel t-tests against chance level (50%). The chance level was subtracted from all reported accuracy values. A statistical threshold of p<0.0001 (uncorrected) at the voxel level, and p<0.05 (family-wise error corrected) at the cluster level was applied, which is sufficient to rule out inflated false-positive rates ^55^. Any regions surpassing this threshold were used as masks for the following decoding analyses (an approach used previously ^16^). The baseline reward decoding is likely partly driven by underlying univariate signal differences, and we do not claim that results reflect differences in response patterns only. This approach does allow us to compare results directly to task-related analyses, which employed the same analysis strategy. The main aim of this analysis was to identify all regions involved in processing reward outcomes. We are not interested in which regions will be found per se, but rather focus on whether reward-related signals will be modified by contingency.

###### Differences in reward outcome coding

Although the baseline decoding analysis should have the maximum power to detect any outcome-related brain regions, results do not allow us to conclude whether outcome processing differed between CR and NCR trials. For this purpose, we performed an additional two searchlight decoding analyses. In the first, we again contrasted high and low reward trials, now only using data from CR trials. In the second, we used only data from NCR trials. If contingent rewards indeed enhance encoding of reward outcomes in the brain, we should see higher accuracies in the CR than in the NCR decoding analysis. Please note, that we only used half the number of trials as before, thus considerably reducing the signal-to-noise ratio in these analyses. We thus expected lower statistical power and smaller effects. Also, whereas the baseline decoding results themselves might be driven by e.g. differences in attentional processing between high and low rewards, looking at *differences* between CR and NCR trials much reduces the impact of any such unspecific differences on decoding results, which are driven more by differences between contingency. We still cannot fully exclude that unspecific processes contribute to these results, however. Lastly, focusing on *differences* between conditions avoids potential issues with double dipping.

###### Similarities in in reward outcome coding

Previous work demonstrated that not all brain regions show a contingency-related modulation of value signals ^10^, and we thus tested whether some brain regions encoded reward outcomes similarly across the contingency conditions. In an additional searchlight decoding analyses, we trained a classifier to discriminate between high and low reward outcomes in the CR condition, and tested its performance in the NCR condition, and vice versa. This resulted in two accuracy maps per subject, which were averaged and then entered into a group analysis just like in the previous analyses. Importantly, only brain regions where the same hyperplane can be used to differentiate neural patterns across both contingency conditions (i.e. in which patterns do not differ substantially) will show above-chance accuracies in this analysis. This so-called cross classification (xclass) analysis provides positive evidence to identify regions that encode reward outcomes independent of the contingency manipulation used here (see also ^26^).

##### Multivariate decoding of tasks

We then employed the same analysis strategy described above to investigate possible effects of outcome contingency on task coding as well. Two GLMs were estimated for each subject, one modelling task-related brain activity at the time of decision-making, and one modelling activity during a subsequent maintenance phase. It has been shown that formation and maintenance of intentions rely on partly dissociable brain networks ^56^, and our design allowed us to estimate independent signals related to both epochs as they were separated by a variable inter-trial-interval.

In the first GLM (GLM_maintenance_), for each of the 5 runs we added regressors for each combination of chosen task (task X, task Y) and reward contingency (CR, NCR). All 4 regressors were locked to the cue onset, the duration was set to cover the whole delay period, during which subjects maintained their task representations. Due to the jittered delay period duration, the modelled signals were dissociated from the task execution and feedback presentation (see also ^17^). These boxcar regressors were then convolved with a canonical haemodynamic response function. A second GLM was estimated (GLM_decisiontime_), in order to extract task-specific brain activity at the time subjects made their choice which of the two tasks to perform. This GLM only differed in the time to which regressors were locked. Although the cue suggested that subjects should make a task choice at that point in time, there is no strong way of controlling the exact point in time at which choices were made in any free-choice paradigm, and they might have been made earlier in principle. It has been shown before that under free choice conditions, subjects choose a task as soon as all necessary information to make a choice is available ^24, 29^. In this experiment, this time point is the feedback presentation of the previous trial, and regressors in this analysis were locked to that event. At this point, subjects can judge whether they e.g. chose the HR or LR task and determine which of the two tasks to perform in the next trial. We used this approach successfully in a previous experiment ^29^, and all further task decoding analyses were performed on both GLMs.

###### Baseline decoding

The task decoding analyses followed the same logic as the reward outcome analyses described above. We first performed a searchlight decoding analysis (radius = 3 voxels, C = 1), contrasting parameter estimates for tasks X and Y in all trials (CR and NCR combined). This analysis has the maximum power to detect any brain regions containing task information, which can be notoriously difficult ^57^. Resulting accuracy maps were normalized, smoothed (6mm FWHM), and entered into a group analysis (t-test vs chance level, 50%). Results were thresholded at p<0.001 (uncorrected) at the voxel level, and p<0.05 (family-wise error corrected) at the cluster level. Again, regions surpassing this threshold were used to define functional regions-of-interest for the following decoding analyses ^16^.

###### Differences in task coding

In order to assess whether task coding is modulated by reward contingency, we repeated the decoding analysis separately for CR and NCR trials. If contingent rewards indeed increase task shielding in the brain, we should see higher accuracies in the CR than in the NCR decoding analysis. This effect should be especially pronounced if both tasks are similar and easily confused, which is the case in our experiment. Again, the power of these analyses is considerably lower than for the baseline analysis.

###### Similarities in task coding

Some previous work suggests that tasks are encoded in a context-independent format in the brain ^17, 18^. Here, we again used cross-classification (xclass) by training a classifier on CR trials and then testing it on NCR trials (and vice versa). Any brain regions showing above chance decoding accuracies in this analysis provides positive evidence of task coding that is similar across contingent vs non-contingent reward conditions. This procedure also ensures that task-related signals are not confounded by potential differences in e.g. cognitive load or expected reward across the CR and NCR conditions, as classifiers are trained and tested only *within* one contingency condition.

###### Control analyses

In order to further corroborate the validity of our results, we performed a number of control analyses. It has been pointed out before, that RT effects might partly explain task decoding results ^25^, although others were unable to show any such effects ^29, 33^. Irrespective of the group-level results of testing for RT differences between contingency conditions or tasks, we decided to conservatively control for RTs effects at the single-subjects level as well. First, we repeated the GLM estimation, only adding reaction times as an additional regressor of non-interest. We then repeated the main decoding analyses, and tested whether accuracy values differed significantly. If RTs indeed explain our task decoding results, we should see a reduction in decoding accuracies when RT effects were regressed out of the data.

Then, we performed a ROI decoding analysis on a brain region that is not related to task-performance in any way, expecting accuracies to be at chance level. We chose the primary auditory cortex for this purpose, defined using the WFU_pickatlas tool (https://www.nitrc.org/frs/?group_id=46, RRID: SCR_007378). If our analyses are not biased towards positive accuracies and are specific to cognitive-control related brain regions, we should see chance level results here.

Additionally, we tested the generalizability of our results in a number of a priori defined ROIs (Supplementary Analysis 4 & 5), assessed whether our decoding procedure was biased towards positive accuracies (Supplementary Analysis 6), and tested whether error rates or potential choice biases affected task decoding results (Supplementary Analysis 7). Exploratory analyses were then performed to assess possible correlations between behavioral measures, questionnaires, and fMRI results (Supplementary Analysis 2).

## Acknowledgements

We would like to thank Anita Tusche, Carlo Reverberi, and Ruth Krebs for valuable discussions on this project. This research was supported by the Research Foundation Flanders (FWO), the European Union’s Horizon 2020 research and innovation program under the Marie Skłodowska-Curie grant agreement No 665501, FWO grant FWO.OPR.2013.0136.01, an ERC StG grant and NWO Vidi grant.

## Author contributions

DW and MB designed the research. DW acquired the data. DW and BF analyzed the data. DW wrote the manuscript. DW, BF, and MB reviewed the manuscript.

## Supplementary Material

### Supplementary Analyses

#### Supplementary Analysis 1

In CR trials, subjects were incentivized to stay in the same task repeatedly, while this was not the case for NCR trials. In order to assess this, we computed the distribution of run lengths for each subject, i.e., the number of trials subjects chose to consecutively perform the same task, separately for CR and NCR trials. If task choices were not based on the random reward outcomes, this distribution can be expected to follow an exponential distribution ^1^. Conversely, the run length for CR trials should be longer than in NCR trials, as participants are expected to repeat a highly rewarding task more often. Differences in run lengths were tested using a one-sided Bayesian t-test.

The average run length in CR trials was 2.54 trials (SEM = 0.08), which was indeed longer than in NCR trials (1.95 trials, SEM = 0.07, BF10 >150, see also Figure Supplementary Analysis 1). Interestingly, we found no evidence for repetition biases in NCR trials, which have been reported before ^1^. Run length in NCR trials was equal to what would be expected if subjects chose tasks randomly (BF10 = 0.20, BF10_r=1.41_ = 0.10), which is necessary, but not sufficient to demonstrate truly random task choices.

**Figure Supplementary Analysis 1.**
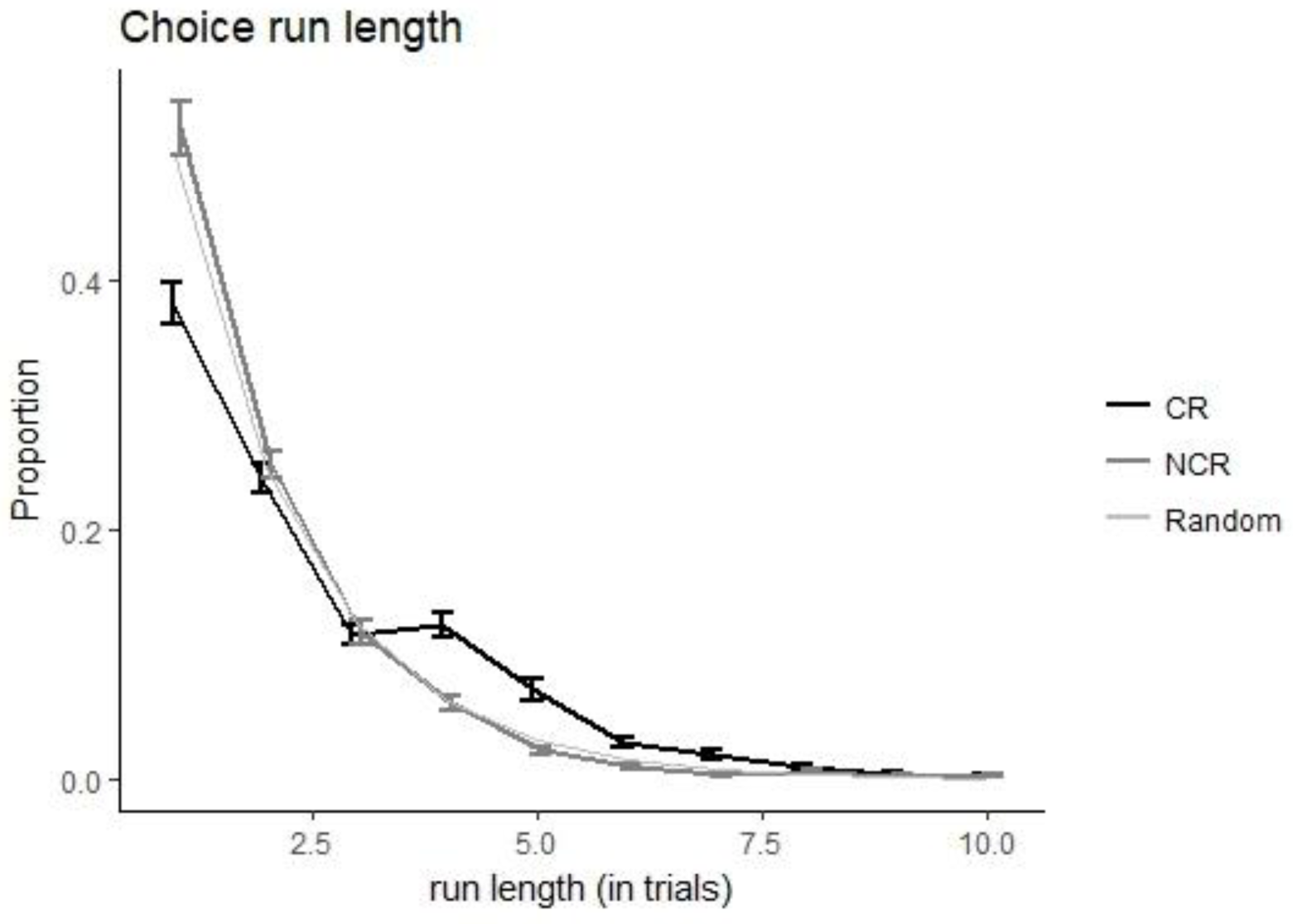
Run length distributions. Depicted are the distributions of run lengths for CR (black) and NCR (dark grey) trials. The expected distribution if choices were random, or at least not guided by reward outcomes, is depicted in light grey.

#### Supplementary Analysis 2

An additional exploratory analysis was performed to correlate performance, questionnaire measures, and decoding accuracies. Several key correlations were assessed using Bayesian correlation analysis (using *bayes.cor.test* form the BayesianFirstAid package) in order to estimate whether they deviated from zero. We report the estimated correlation (r), the probability of the correlation being above or below zero (p_r>0_, p_r<0_), and the 95% credible intervals (95% CI). If the interval does not contain zero, the correlation is larger or smaller than zero.

##### Correlating performance with decoding results

First, we assessed whether performance in the tasks was correlated with decoding results. We only found successful performance in CR trials to be correlated with the degree of reward signal amplification in CR trials (as compared to NCR trials), r = 0.52, p_r>0_ > 0.99, 95% CI = [0.26, 0.76]. The more reward signals were amplified, the more successful subjects were in performing the reversal learning task, which demonstrates the behavioral relevance of the reward signal amplification.

##### Correlating performance with questionnaire results

We found success in CR trials to be negatively correlated with motor impulsivity, r = −0.37, pr<0 = 0.98, 95% CI = [-0.66,-0.08]. The relation to impulsivity was specific to motor impulsivity, we found no strong relation of CR performance with either attentional or non-planning impulsivity. This finding suggests that (motor) impulsive subjects were worse in performing the reversal-learning task (see also ^2^). We found no correlation of success with either sensitivity to reward (r = 0.11, p_r>0_ = 0.74, 95% CI = [-0.23, 0.44]), or the need for cognition (r =0.20, p_r>0_ = 0.87, 95% CI = [-0.12, 0.52]), despite the fact the need for cognition has previously been associated with reward decision-making ^3^.

##### Correlating decoding with questionnaire results

We found task decoding accuracies in the dmPFC at the time of decision-making to be positively correlated with non-planning impulsivity, r = 0.33, p_r>0_ = 0.92, 95% CI = [0.018, 0.63], suggesting that impulsive subjects had stronger task representations right after all necessary information was presented to make a choice. Non-planning impulsivity has been linked to reward decision-making on the behavioral level previously ^2^, but this result suggests that a similar relation might be present at the neural level as well. Overall, the relation of impulsivity and performance / decoding results was unexpected, and future research should be targeted more at explaining how impulsivity affect the neural basis of reward decision-making.

**Figure Supplementary Analysis 5:**
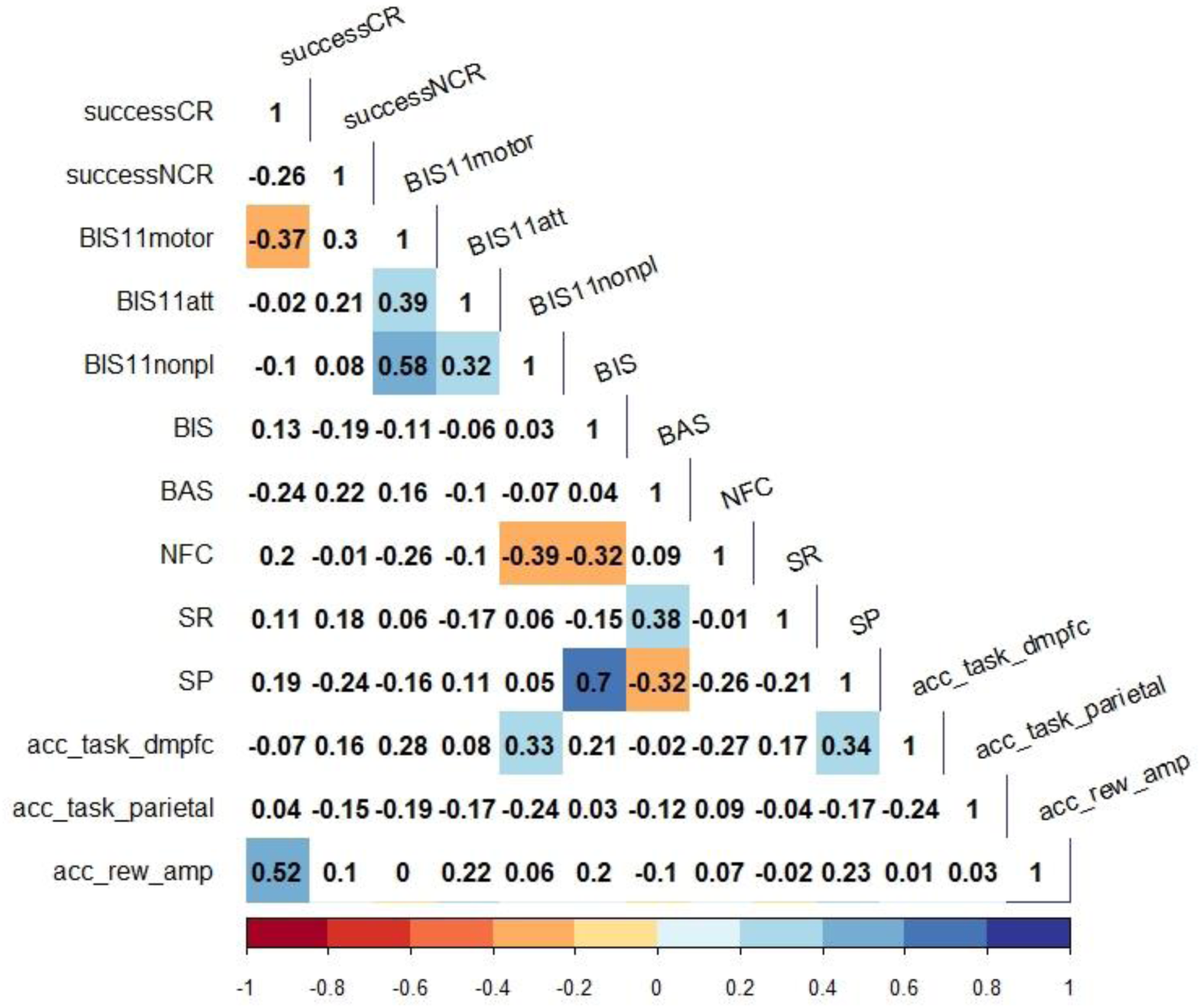
Correlation analysis. Depicted are pairwise correlations (estimated using *bayes.cor.test* form the BayesianFirstAid package) of: % high reward choices in CR trials (successCR), % high reward outcomes in NCR trials (successNCR), motor impulsivity (BIS11motor), attentional impulsivity (BIS11att), non-planning impulsivity (BIS11nonpl), behavioral inhibition (BIS), behavioral approach (BAS), need for cognition (NFC), sensitivity to reward (SR), sensitivity to punishment (SP), decoding accuracies in the baseline task decoding analysis in the dorso-medial PFC (acc_task_dmpfc) and parietal cortex (acc_task_parietal), and an index of reward signal amplification (acc_rew_amp). This index was computed by subtracting accuracy values from the reward outcome decoding in NCR trials, from the accuracy values in CR trials. Only regions that showed successful cross-classification of reward signals across contingency conditions were included here. This procedure leads to a global measure of how strongly reward signals were amplified in the CR condition, as compared to the NCR condition. The plot was generated using the *corrplot* package in R. Number show the correlation coefficients. Correlations in white cells did not differ from zero (i.e. the 95% CI included zero), correlations in colored cells did.

#### Supplementary Analysis 3

Given that decoding tasks separately in CR and NCR trials has considerably less power than decoding tasks in both CR and NCR trials together, we performed an additional control analysis to determine if the reduced power explains the absence of any differences between task coding in CR and NCR trials.

Previous research demonstrated that task signals in the brain are modulated by associated reward outcome magnitude, i.e. tasks that lead to a high reward show stronger coding than tasks that lead to no reward ^4^. Based on this, we expected to find a similar effect in our experiment as well. We tested whether tasks directly following a HR outcome were encoded more strongly than tasks directly following a LR outcome. For this purpose, we estimated another set of GLMs for each subject, only now splitting trials into following HR vs LR (instead of CR vs NCR trials). In all other respects, this analysis was identical to the task decoding presented in the main body of the paper. We then extracted decoding accuracies from the parietal, right aMFG, and dmPFC clusters used in the main analysis, separately for HR task decoding and LR task decoding. Using Bayesian t-tests vs chance and a paired Bayesian t-test, we assessed whether there were any differences between these accuracies. We expected task coding to be stronger in HR trials especially at the time of decision making, as this time point is closer to the reward feedback presentation than the maintenance period.

At the time of decision making, we found strong task coding in the dmPFC in HR trials (60.67%, BF10 > 150), but no task coding in LR trials (51.07%, BF10 = 0.33). Crucially, there was strong evidence for a difference between these values (BF10 = 31.89). During the maintenance period, the parietal cortex showed task coding in the HR condition (54.95%, BF10 = 5.83), but not in the LR condition (52.32%, BF10 =0.59). We found no evidence for any difference between these values however (BF10 = 0.66). In the right aMFG a similar picture emerged. We found task coding in HR trials (56.07%, BF10 = 14.36), but not in LR trials (52.82%, BF10 =1.51), and no difference between these values (BF10 = 1.03).

Thus, we find evidence for differences in task coding, although the evidence is stronger at the time of decision-making than the maintenance period. This demonstrates that our analysis approach can in principle detect differences in task coding in the current data-set, yet we still fail to find any such differences between contingent and non-contingent trials.

#### Supplementary Analysis 4

We assessed task information in a number of a priori defined ROIs. First, we attempted to replicate results from some previous experiments ^5, 6^. Both of these experiments assessed whether task coding is modulated by external variables. While Wisniewski et al. (2016) looked into effects of free choice on task coding, Loose et al. (2017) looked into effects of cognitive control demands on task coding. Both studies found a fronto-parietal network to encode task sets and found them to be independent of the experimental manipulation employed. Given that the tasks employed in these studies were different from the one used here (e.g. addition vs subtraction of two numbers), a replication of these findings in the current dataset would show a strong generalizability of our results. We thus extracted functional ROIs from both papers (Wisniewski et al. 2016: left parietal cortex, left PFC, Brodman area 8; Loose et al. 2017: left parietal cortex, left PFC), and extracted accuracy values for all voxels within the ROI for all four decoding analyses performed (baseline, CR, NCR, xclass), which were then averaged. One-sided Bayesian t-tests across subjects were performed to assess whether they were above chance.

We were able to replicate Loose and colleagues’ left parietal results (baseline BF10 = 133.69; CR BF10 = 0.68; NCR BF10 = 0.54; xclass BF10 = 33.17). Although somewhat weaker, we also replicated their right parietal results (baseline BF10 = 8.49; CR BF10 = 0.77; NCR BF10 = 0.14; xclass BF10 = 8.10). However, we were unable to detect task information in left PFC (baseline BF10 = 0.49; CR BF10 = 0.21; NCR BF10 = 0.44; xclass BF10 = 0.29), which is in line with the original paper, where PFC findings were also somewhat less robust. Additionally, we were able to replicate Wisniewski and colleagues’ left parietal finding (baseline BF10 > 150; CR BF10 = 0.80; NCR BF10 = 0.47; xclass BF10 = 87.28), as well as left BA8 (baseline BF10 = 9.3; CR BF10 = 0.39; NCR BF10 = 0.36; xclass BF10 = 3.09), but not the left PFC (baseline BF10 = 0.59; CR BF10 = 0.37; NCR BF10 = 0.16; xclass BF10 = 0.38). Thus, we find task coding to be contingency-independent even in ROIs that were defined using independent datasets. We further show that task information is most consistently found in the parietal cortex, but less so in prefrontal cortex.

#### Supplementary analysis 5

Some previous work suggested that task information can be found in the multiple demand (MD) network ^7^, and that task coding in this network changes flexibly with changing task demands ^8^. In order to test whether this was also the case in our dataset, we extracted accuracy values for all four decoding analyses (from bilateral functional MD ROIs (provided by ^9^), specifically the anterior insula (aINS), cerebellum, inferior frontal gyrus pars opercularis (IFGop), intraparietal sulcus (IPS), middle frontal gyrus (MFG), pre-central gyrus (precG), supplementary and pre-supplementary motor area (SMA/preSMA), as well as thalamus. We then tested whether decoding accuracies were higher in CR than in NCR trials, and/or whether task coding was contingency-independent in these regions. Averaging across all MD regions, we found strong evidence for the presence of task information (52.23%, SEM = 0.61%, BF10 = 69.08, see Figure below). We found no evidence for a higher accuracy in CR, as compared to NCR trials (BF10 = 0.37). Furthermore, we found task coding to be contingency-independent (52.02%, SEM = 0.67%, BF10 = 14.52). Accuracies in the baseline and cross-classification analysis did not differ (BF10 = 0.19). Looking at individual MD regions, we found task information in the aINS (52.25%, SEM = 1.00%, BF10 = 3.23), IPS (52.83%, SEM = 0.72%, BF10 = 131.02), MFG (52.44%, SEM = 0.90%, BF10 = 8.26), precentral gyrus (52.48%, SEM = 0.87, BF10 = 9.86), but not in the cerebellum (50.85%, SEM = 0.90%, BF10 = 0.44), IFGop (52.11%, SEM = 1.02%, BF10 = 2.31), SMA/preSMA (51.48%, SEM = 1.07%, BF10 = 0.77), and thalamus (50.58%, SEM = 1.06%, BF10 = 0.29). None of these regions showed a higher accuracy in CR than in NCR trials (BFs10 <= 0.60). However, in all of those regions the accuracy in the baseline and xclass analyses was equal (BFs10 <= 0.28). This suggests that the MD network encodes tasks in a contingency-independent fashion, and shows that the current context does not affect task coding in the MD network. Also, not all parts of the MD network seemed to be encoding tasks in our experiment.

**Figure Supplementary Analysis 2.**
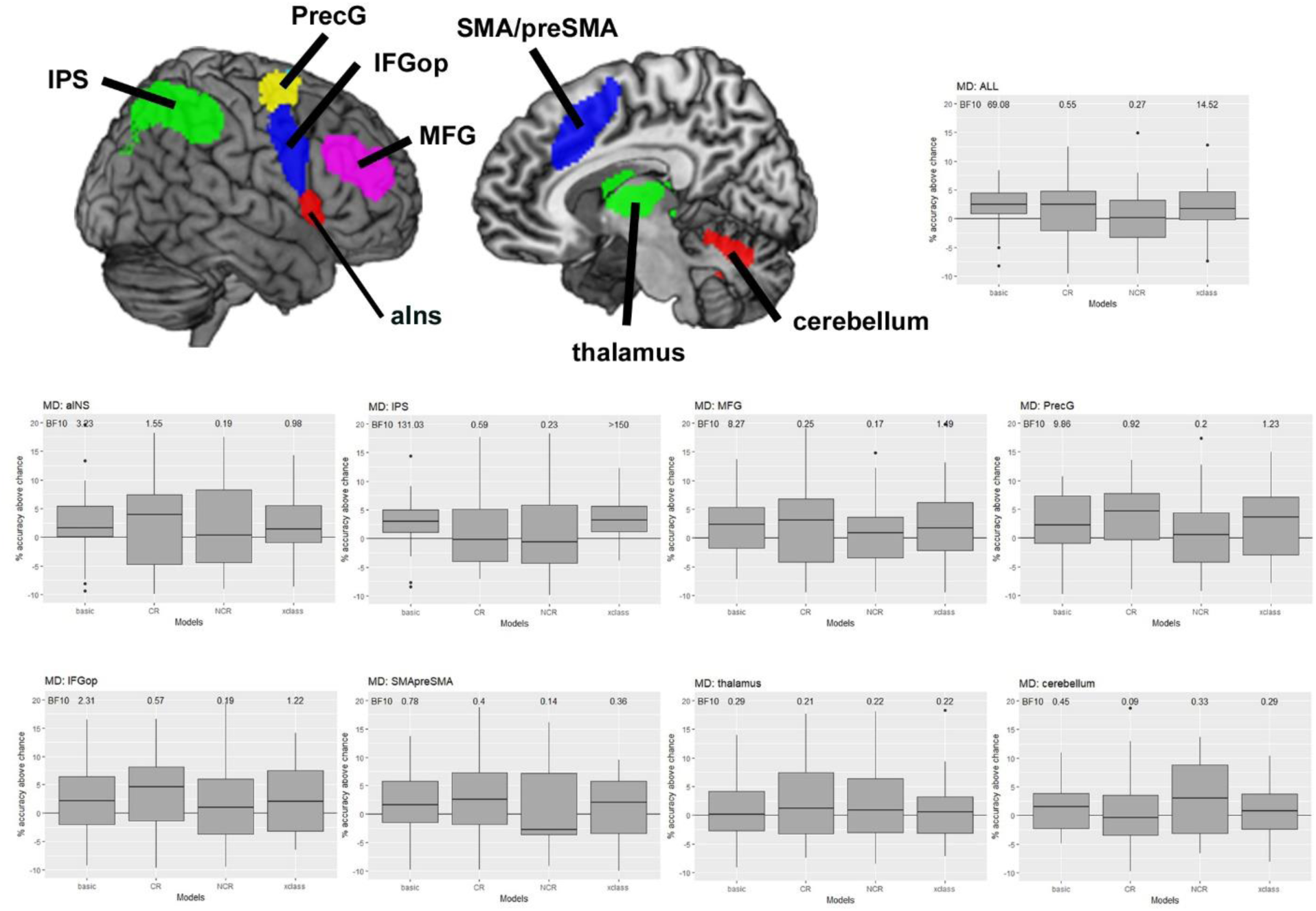
Task information in the multiple demand (MD) network. Depicted are task decoding results in the bilateral functional ROIS provided by Fedorenko, Duncan, & Kanwisher (2013), specifically the anterior insula (aINS), cerebellum, inferior frontal gyrus pars opercularis (IFGop), intraparietal sulcus (IPS), middle frontal gyrus (MFG), pre-central gyrus (precG), supplementary and pre-supplementary motor area (SMA/preSMA), as well as thalamus.

#### Supplementary analysis 6

In order to assess whether our decoding procedure was biased towards positive accuracy values, we empirically estimated the chance level and tested if it was indeed 50% as we assumed. For this purpose, we performed a permutation analysis (n = 1000 permutations per subject, as implemented in the Decoding Toolbox, using the same regressors and contrasts as the baseline task decoding analysis) in order to estimate the null distribution of our data. We took the mean of the null distribution as our empirical estimate of the chance level, and tested whether it deviated from 50% (using a two-sided Bayesian t-test). If there were some global biases in our decoding procedure, chance level should deviate from 50%. The estimated chance level was 49.98%, which did not differ from 50% (BF01 > 150). Thus, comparing our decoding accuracies against a chance level of 50% was valid.

#### Supplementary analysis 7

Although overall error rates were very low, and we found no evidence for persistent choice biases across our sample, there might still be individual subjects that do show e.g. high error rates, which might affect our task decoding results by decreasing signal-to-noise ratio. Although the effect of few outlier subjects should be small given our large sample size, we still chose to conservatively control for such (unlikely) effects. We first excluded subjects with the highest error rates (more than 1.5*IQR above average, i.e. error rate > 13.92%), and then excluded subjects with the strongest choice biases (more than 1.5*IQR above average, i.e. percent task X choices > 61.99% or < 38.73%). We then tested whether each regressor in the task decoding analysis in all remaining subjects could be estimated from at least 6 trials. If a regressor could only be estimated from fewer trials, that run was excluded from the analysis due to the low signal-to-noise-ratio. Subjects in which more than 1 run was thusly excluded were altogether excluded from the analysis. These criteria were highly similar to the criterion used in ^10^, which proved an effective control. After excluding these subjects, we repeated the main four analyses (baseline, CR, NCR, xclass) on the remaining subjects and tested whether they differed from the analysis including all subjects.

Using these highly conservative exclusion criteria, we removed 2 subjects due to their error rate, 5 subjects due to their choice biases, and 4 subjects due to the small number of trials, leading to a sample size of 24 subjects. Even though statistical power was considerably lower because of the smaller sample size, we were still able to detect task information in the parietal cortex (55.09%, SEM = 0.78%, BF10 >150), which was again reward-independent (53.63%, SEM = 0.98%, BF10 = 60.25), and the same was true for the aMFG (54.74%, SEM = 1.09%, BF10 >150, and 53.53%, SEM = 1.35%, BF10 = 6.57, respectively). These results are even numerically larger than in the original analysis, and neither error rates not choice biases were found to affect the reported task decoding results.

**Supplementary Figure 1.**
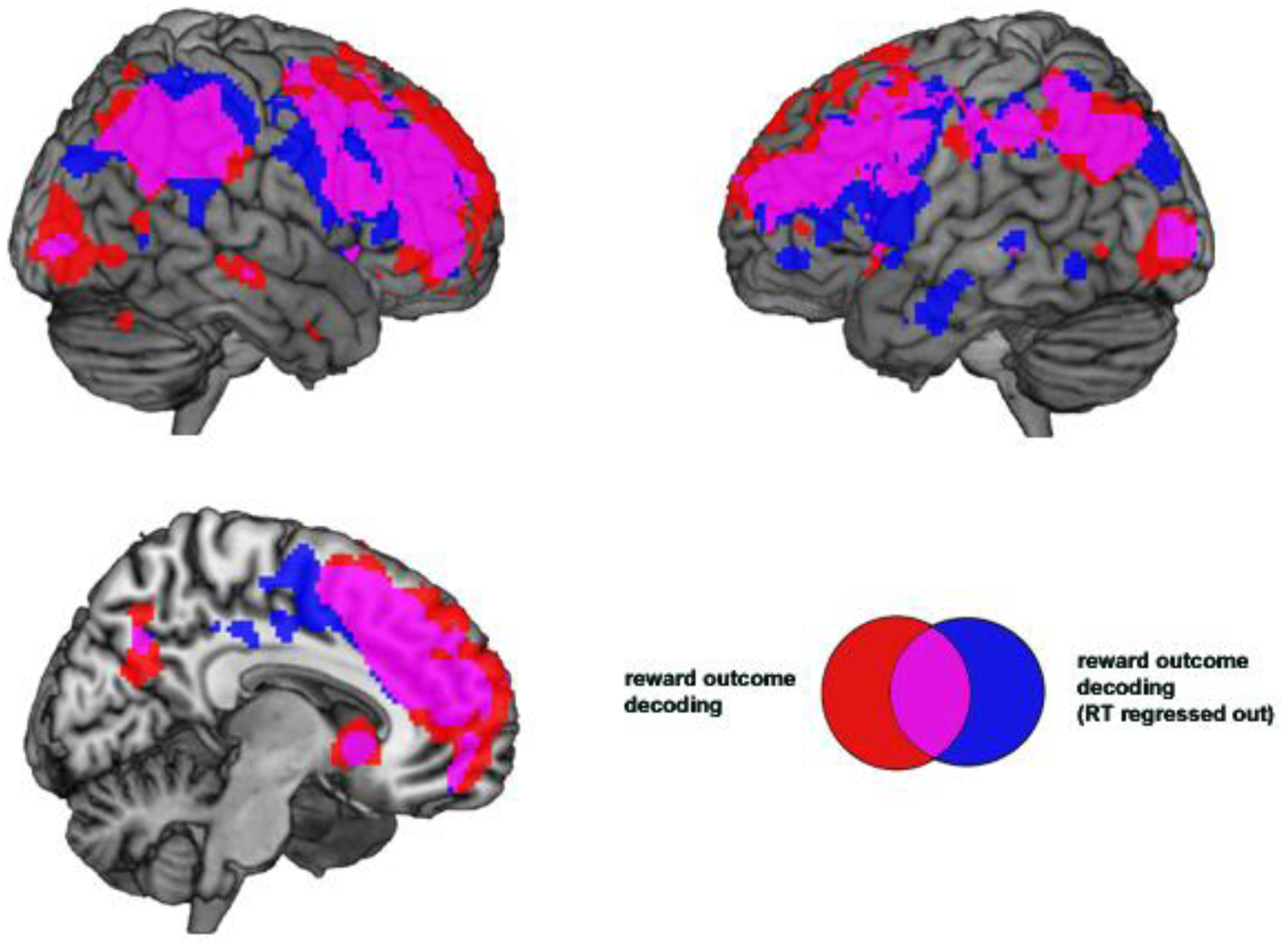
Controlling RT-effects in reward outcome decoding. We repeated the reward outcome decoding analysis, using a similar first-level GLM to estimate signals (4 regressors: high contingent reward, low contingent reward, high non-contingent reward, low non-contingent reward, all locked to feedback onset). Additionally, we now added parametric regressors of non-interest capturing RT-related variance in the data. The rest of the analysis was identical to the reward outcome decoding analysis presented in the main body of the text. Results from the reward outcome decoding analysis (red), and the same analysis with RT-related effects regressed out of the data (blue) are depicted. As can be seen, the overlap (magenta) between both analyses is substantial. Results depicted at p < 0.05 (FWE, corrected at the voxel level). This indicates that controlling for RT did not strongly alter our results.

